# Evolutionary Analyses of RNA Editing and Amino Acid Recoding in Cephalopods

**DOI:** 10.1101/714394

**Authors:** Mingye (Christina) Wang, Erik Mohlhenrich

## Abstract

RNA editing is a post-transcriptional modification process that alters nucleotides of mRNA and consequently the amino acids of the translated protein without changing the original DNA sequence. In human and other mammals, amino acid recoding from RNA editing is rare, and most edits are non-adaptive and provide no fitness advantage (*1*). However, recently it was discovered that in soft-bodied cephalopods, which are exceptionally intelligent and include squid, octopus, and cuttlefish, RNA editing is widespread and positively selected (*2*). To examine the effects of RNA editing on individual genes, we developed a “diversity score” system that quantitatively assesses the amount of diversity generated in each gene, incorporating combinatorial diversity and the radicalness of amino acid changes. Using this metric, we compiled a list of top 100 genes across the cephalopod species that are most diversified by RNA editing. This list of candidate genes provides directions for future research into the specific functional impact of RNA editing in terms of protein structure and cellular function on individual proteins. Additionally, considering the connection of RNA editing to the nervous system, and the exceptional intelligence of cephalopod, the candidate genes may shed light to the molecular development of behavioral complexity and intelligence. To further investigate global influences of RNA editing on the transcriptome, we investigated changes in nucleotide composition and codon usage biases in edited genes and coleoid transcriptome in general. Results show that these features indeed correlate with editing and may correspond to causes or effects of RNA editing. In addition, we characterized the unusual RNA editing in cephalopods by analyzing ratio of radical to conservative amino acid substitutions (R/C) and distribution of amino acid recoding from editing. Our results show that compared to model organisms, editing in cephalopods have significantly decreased R/C ratio and distinct distribution of amino acid substitutions that favor conservative over radical changes, indicating selection at the amino acid level and providing a potential mechanism for the evolution of widespread RNA editing in cephalopods.

## Introduction

According to the central dogma, DNA is transcribed into messenger RNA, which is translated into protein. In this process, there is a direct correspondence between the DNA sequence and the amino acid sequence. One exception to the central dogma is direct RNA editing, which alters the nucleotides of mRNA and potentially the amino acids of the translated protein without changing the original DNA sequence. A-to-I editing (herein simply referred to as RNA editing) is the most common type of RNA editing in vertebrates, in which adenosine is deaminated to form inosine, which behaves as guanine in translation (*3*). This process is carried out by the enzyme ADAR (Adenosine Deaminase Acting on RNA), which binds to double-stranded RNA with preferred sequences. Overall, RNA editing is rare in mammals. In humans, only about 3% of mRNA are edited, of which only 25 editing sites are conserved across mammals (*2*). Studies have shown that the vast majority of edits in human are non-adaptive and likely the result of promiscuous editing by ADAR (*4*). Furthermore, most editing occurs in non-coding transcripts encoding Alu repeat elements.

In contrast to mammals, a 2017 study demonstrated that mRNA is extensively edited in coleoids, the group of soft-bodied cephalopods that include squid, octopus, and cuttlefish (*2*). In *Octopus bimaculoides*, more than 900,000 editing sites were found by analyzing RNA from 12 different tissues, of which 12% are in coding regions. Furthermore, 65% of these editing sites in coding region are non-synonymous (resulting in an amino acid substitution, known as protein recoding), and conserved sites tend to have higher editing levels and a higher proportion of non-synonymous substitutions. In total, 1146 editing sites were found to be conserved over all four coleoid species studied (*Octopus vulgaris, Octopus bimaculoides, Doryteuthis pealeii, Sepia oficianalis*), which diverged 350-480 mya, in sharp contrast with the 25 transcripts conserved across mammals, which diverged only 150-200 mya (*5*).

Previous works have shown that RNA editing is crucial in regulating the nervous system in mammals. Mammalian ADARs have the highest expression level in the central nervous system (*1*). Although protein recoding by RNA editing in mammals is rare, several of the few recoding editing events involve proteins related to the nervous system, including glutamate receptor subunit GluR2 and potassium channel Kv1.1. Deletion of ADAR2 in mice, for example, cause frequent epileptic seizures due to neuronal death, and the effect seems to be due to the lack of only one essential edit (*6*). The connection between RNA editing nervous system is also discovered in cephalopods and is even more pronounced (*2*). Protein recoding (editing at non-synonymous sites) from RNA editing in coleoids is most significant in the nervous system. In *Octopus bimaculoides*, editing in neural tissues is 2-fold higher and protein recoding events are more frequent. In addition, the closely related nautilus species, which have simpler brains and are behaviorally less complex than coleoids, have significantly fewer RNA editing events, while coleoids with extensive RNA editing have the largest nervous system in invertebrates and are known for their exceptional intelligence (*2*). Together, these discoveries suggest an important role for protein recoding by RNA editing in neural and behavioral complexity.

To better understand the effects of RNA editing in cephalopods, we developed a quantitative metric to identify edited genes that are most impacted by RNA editing. One benefit of RNA editing over regular DNA mutations is the flexibility it allows for expression of different versions of a transcript at different time periods, potentially in response to shifting environmental conditions, thus offering another level of temporal regulatory control (*6*). Another major benefit of RNA editing is the diversity it creates: since not all mRNA transcripts may be edited, this allows different versions of the same sequence (both edited and unedited) to be expressed in the same cells (*2*). In fact, for genes with multiple editing sites, if the combinatorial editing is realized, the number of different variants for each encoded protein would increase exponentially as the number of editing sites within a gene increases (*7*). Therefore, we aimed to quantify this diversity and use it as the main evaluation for identifying candidate genes by developing a quantitative measure we named “diversity score” that ranks all genes containing at least one editing site by the level of protein diversity generated by RNA editing. The main factors that affect the amount of molecular diversity include the editing level and the specific amino acid change that is caused by the edited site. Editing level measures the relative amount of RNA transcripts that are edited, and vary from nearly 0% to 100%. A value near 50% achieves maximal diversity, as both types of transcripts are expressed evenly inside the cells, similar to the concept of species evenness for ecological systems (e.g. in Shannon diversity index), and is thus taken into account when assessing diversity (*8*). For the amino acid change, two aspects were considered in the diversity score. Chemically, amino acid substitutions between two dissimilar amino acids (i.e. a radical substitution, such as when a positively charged amino acid is substituted for a negatively charged amino acid) are more likely to impact the proteins significantly in structure and function than substitutions between two chemically similar amino acids (i.e. a conservative substitution), generating higher diversity (*9*). Therefore, chemical properties of the amino acid change caused by an editing site are taken into account in the diversity score.

Although physicochemically radical substitutions are more likely to impact protein structure and function on average, this may not be true at all residues in a protein. Conversely, the constraints on a particular residue may cause even conservative substitutions to have substantial effects. For example, a conservative substitution from arginine to histidine in the ATPase ATP7A causes Menkes disease, a fatal illness in infants (*10*). The constraints at a specific residue can be accounted for by incorporating evolutionary information - if having a certain amino acid at a site has a damaging effect on the protein structure and function, then it is expected that the particular amino acid would be very rare across matching protein sequences of numerous species, as natural selection tends to eliminate deleterious substitutions (*11*).

Here, we incorporate editing level with measures of evolutionary and physicochemical properties of amino acid substitutions to develop a protein diversity score that quantitatively assesses the amount of combinatorial diversity generated by RNA editing. Using this metric, we calculate diversity scores for each gene in the four cephalopod species and compile a list of top 100 genes across the cephalopods. We further analyze this list of candidate genes with gene ontology enrichment, disease annotations, protein-protein interaction networks, and comparisons to genes edited in model organisms. These results highlight connections between RNA editing and nervous system development, and provide a list of candidate genes for further studies into the role of RNA editing in neural and behavioral complexity in cephalopods.

In addition to specific molecular effects, widespread RNA editing may also lead to global changes in the transcriptome. For example, given that RNA editing in cephalopods acts on adenosine, one might expect the genome content of adenosine is increased relative to related species. Previously it has been shown that regions around editing sites (up to 100 nucleotides on each side) display elevated GC content, which facilitate formation of stable double-stranded regions required by ADAR enzymes to bind and initiate editing *(2*). We therefore might expect increases in overall GC content in cephalopod species. Besides nucleotide composition, another potential consequence of widespread RNA editing is changes in codon usage bias, in which certain codons are used at higher frequencies than other synonymous codons (*12*). Codon usage bias has various influences on processes related to gene expression--for example, codon usage of highly expressed genes is selected to increase translation efficiency (*13*). Within the context of RNA editing in cephalopods, given synonymous codons that code for the identical amino acid, we might expect the codon containing a “A” to be preferred since it enables further editing that may diversify the protein. Conversely, the A-containing codon may also be actively avoided at crucial positions of a gene to prevent any editing. These changes in nucleotide composition and codon usage biases may affect many other fundamental processes, for example mutational biases and translation efficiency, adding to the cost-benefit trade-off of widespread RNA editing, in addition to the already discovered reduction of polymorphisms caused by the maintenance of necessary structures surrounding editing sites (*2*).

One major question arising from the original discovery of widespread RNA editing in cephalopods is the cause of its evolution in cephalopods and not other species. We hypothesize that overall genomic characteristics may play a part--initial changes in cephalopod genome may have facilitated RNA editing, which in turn led to more biases towards genomic content that favors RNA editing. Investigation into nucleotide composition and codon usage bias in cephalopods would thus not only address the effects of widespread RNA editing on global transcriptome characteristics, but also help elucidate the factors that led to the evolution of the process itself.

Therefore, we examined and compared nucleotide content and codon usage bias in coleoids with nautilus and aplysia. By building predictive models of editing, we show that nucleotide composition and codon usages indeed contain information on editing. Further analysis showed that contrary to expectation, edited genes do not appear to have different nucleotide content (specifically proportion of adenosine and GC) than non-edited genes, while coleoids in general do have higher levels of A, but lower levels of GC compared to nautilus and aplysia. Meanwhile, several codons show biases that follow expected trends from RNA editing and may warrant further attention.

Lastly, we analyzed the amino acid recoding from RNA editing to gain insights into the strength of natural selection acting on RNA editing and the potential causes of widespread editing. Previously it has already been shown that nonsynonymous edits are suppressed in mammals, one evidence that the majority of editing in mammals is non-adaptive and not under selective pressure, likely results of promiscuous editing by the ADAR enzyme (*4*). In cephalopods, nonsynonymous-to-synonymous ratio (N/S) is neutral overall, but the ratio increases as editing levels increase, indicating positive selection for highly edited sites (*2*). These N/S analysis are focused on the nucleotide level, and therefore we investigated whether if similar pressures act on the amino acid level by calculating the ratio of radical-to-conservative amino acid substitutions (R/C). A lower R/C ratio indicates stronger purifying selection, since radical substitutions are more likely to be deleterious and actively removed when selection is strong (*9*). Our results show that cephalopods editing have lower R/C ratios compared to human, mice, and fly, indicating strong purifying selection in addition to the positive selection previously discovered. To further characterize the amino acid recoding, we examined the distribution of amino acid substitutions by RNA editing, and cephalopods show a distinct pattern that more strongly bias conservative substitutions over radical substitutions, compared to patterns in human, mice, and fly. These results provide insights into the amino acid recoding from RNA editing and the selection pressure acting on it, and offer potential clues to the evolution of widespread RNA editing in cephalopods.

## Methods

### I. Transcriptome and Editing Data Acquisition

Transcriptome and editing data of four coleoid species (*Octopus vulgaris, Octopus bimaculoides, Doryteuthis pealeii, and Sepia oficianalis*; hereafter abbreviated as oct vul, oct bim, squid, and sepia), and transcriptome data of a nautiloid (*Nautilus pompilius*; hereafter referred to as nautilus) and a gastroid mollusk (*Aplysia californica*; hereafter referred to as aplysia) were acquired from Liscovitch-Brauer et al., 2017 (see Supp. I for a sample section of the dataset). Trancriptome data of the four coleoid species were modified to reflect unedited sequences by matching editing sites with editing level greater or equal to 50% to the transcriptome nucleotide and mutating it from G to A. In total, 117842 sites in oct vul, 76862 sites in oct bim, 82975 editing sites in squid, and 130636 sites in sepia were used. Editing data of mouse (mm9) and fly (dm3) were acquired from the Rigorously Annotated Database of A-to-I RNA Editing (RADAR) (*14*). Editing data of human was acquired from both RADAR and REDIportal from the GTEx project (*15*).

### II. General Data Processing

Data cleaning, transformations, and calculations were performed using the online bioinformatics platform Galaxy (*16*) and other tools including RStudio, Unix shell, and Microsoft Excel. The software package EMBOSS was used for sequence translation (*17*), and the software package CAT was used for nucleotide composition and codon usage analysis (*12*).

### III. Diversity Score Calculation

The diversity score is composed of the editing level score and the amino acid score. Diversity score of a gene is calculated as the product of the editing level score and the amino acid score summed over all editing sites in the gene.

Editing level score of each editing site is calculated as to reflect the larger diversity achieved with editing values close to 50%:

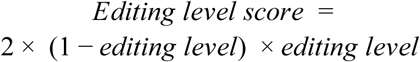

In this way, the diversity score of one gene represents the mean distance, measured by the sum of amino acid scores per editing site, between two randomly selected transcripts, each containing a portion of the amino acids substituted, with distribution as according to the editing level. Two issues arising from data limitation should be noted: first, the distribution used in this method assumes all editing sites to be independent, while in reality not all combinatorial diversity may be achieved in the actual cellular environment: previous studies have shown that certain editing sites are correlated (*18*), and therefore some transcripts may not be realized or occur at a lower frequency than expected. Second, the editing level data was pooled over multiple tissues, and therefore it is possible that only certain sites are edited in one tissue, so the actual diversity is lower than expected. Thus, the diversity score calculated here represents only a theoretical maximum diversity that may be achieved.

Two different amino acid scores were calculated, one focusing on the general chemical effect of the substitution, and the other incorporating site-specific evolutionary information. The universal evolutionary index (UEI) was used as a proxy for the physicochemical significance of an amino acid substitution (*19*). UEI was derived from comparisons of sequences across a large number of proteins in various species, and describes the likelihood that one amino acid is substituted for another across evolutionary time. Since more chemically radical substitutions are on average more deleterious than conservative substitutions and more likely to be removed by natural selection (*9*), UEI scores serve as an indication of the general radicalness of an amino acid substitution chemically. The other amino acid score utilizes PSSM scores (Position Specific Scoring Matrix) from delta-BLAST, which searches in a database of sequences from various species to find sequences that are similar to the edited protein in cephalopods (*20*). PSSM is position specific, providing different scores for the same amino acid substitution at different positions. Higher scores correspond to more frequent amino acids (i.e. many species have the particular amino acid at the particular site of the protein) and often indicate critical functional residues. Editing that produces two amino acids with highly disparate PSSM scores (i.e. one amino acid has a very high score, indicating it is commonly found at this location in other species, while the other amino acid has a low score, indicating it is rarely found) indicates the coexistence of a highly conserved, canonical form of the protein, as well as a novel form, and thus was taken to be highly diverse in the diversity score.

Numerically, the UEI score is defined as the reciprocal of the UEI value because in the original formulation of the UEI, the higher the value, the more conservative or chemically similar an amino acid change. The PSSM score is defined as the absolute value of the difference between the scores for the edited amino acid and the unedited version. PSSM values were calculated by searching cephalopod protein sequences against the CDD delta database with E-value = 1e-5 as the threshold for matches. Diversity score-UEI is defined as *editing level score* × *UEI score*, summed over all editing sites per gene. Similarly, diversity score-PSSM is defined as *editing level score* × *PSSM score* summed over all sites. An additional measure we named “novelty score” was calculated as the product of the absolute editing level and the directional difference of the PSSM scores (i.e. PSSM score of the edited amino acid subtracted by the score of the original amino acid, taken the negative to account for the sign, and if negative is replaced by 0 since an edit should not diminish novelty) summed over all sites. This score identifies sites that approximate a novel mutation, where the high editing level indicates strong selection for the edited version of the protein, and the large directional difference in the PSSM score (i.e. the unedited amino acid has a high score and the edited amino acid has a low score) indicates the edit transforms a common, canonical form of the protein into a rare, novel form.

Using these measures, the diversity scores (UEI, PSSM, and novelty) of all edited genes in each of the 4 cephalopod species were calculated. The scores are combined across the 4 species: the genes are matched across the cephalopod species by using their matches to human genes (Swiss Prot) and the diversity scores from each species were averaged. Due to the existence of paralogs, multiple genes in one cephalopod species may be mapped to a single human gene. In this case, the mean score of all possible combinations was taken. Since UEI and PSSM diversity scores differ in numerical scales, these two scores are scaled by each measure using the maximum UEI or PSSM score in the entire data set and summed together to generate an overall ranking of edited genes (which we refer to as the “overall” gene list). In addition, novelty scores of individual editing sites in each species, as well as conserved sites across all 4 species, were used to rank the sites for specific editing site candidates.

### IV. Functional Analyses of Edited Genes

The online database STRING was used to generate protein-protein association networks of candidate genes (*21*). Experiments, databases, co-expression, neighborhood, gene fusion, and co-occurrence were used as sources of interactions, and medium confidence (0.400) was used for the minimum required interaction score. Interaction networks were generated for the top 100 and 200 candidate genes overall and were then analyzed to identify highly connected genes we call “hub” genes. Specifically, genes that connected to at least 5 genes in the top 100 list, and at least 10 in the top 200 list, were highlighted as hub genes.

A set of genes associated with human intelligence was acquired from a GWAS study (*22*). This intelligence gene set was compared to genes edited in cephalopods, and candidate genes that are included in this gene set are highlighted.

Gene ontology enrichment results were calculated using the online tool GOrilla (*23*) with a single list of genes ranked by diversity score, with p-value threshold = 1E-3.

Disease annotations for edited genes were acquired from the curated gene-disease associations dataset on DisGeNET (*24*). A cutoff of 0.6 was used for filtering out weak associations. Disease enrichment was calculated using the minimum hypergeometric score method used in gene ontology enrichment calculations in GOrilla with a single list of genes ranked by diversity score (*23*). Diseases associated with fewer than 10 genes in the entire dataset were removed.

For analyses requiring gene names, UniProt IDs were mapped to gene names using the UniProt website (https://www.uniprot.org/mapping/).

### V. Codon Usage Bias Analysis

Nucleotide composition and codon usage analysis were performed on the four coleoid species (oct vul, oct bim, squid, and sepia) and two outgroups (nautilus, aplysia) using the software package CAT with translated transcriptome data (*12*). Edited genes from the four coleoid species were extracted and separate codon usage bias values were calculated.

To evaluate the hypothesis that codon usage bias profiles of edited genes are distinct from unedited genes, machine learning algorithms were used to predict a) whether if a gene is edited and b) cumulative editing level normalized by length, using observed levels of all 61 codons (excluding 3 stop codons), nucleotide composition (GC and AG proportions overall and at positions 1, 2, 3 in codon), and codon deviation coefficient (CDC), an overall measure of codon usage bias calculated by CAT, for a total of 70 features. For both tasks, transcriptome data from only *Octopus bimaculoides* were used to avoid potential interference from global differences between species. The data was split into 75% for training and 25% for testing. For classifying if a gene is edited, a simple logistic regression model (R package glm) was enough to achieve substantial accuracy, while for predicting editing level, random forest was used (R package randomForest). Both are implemented through the R package caret (for random forest, RMSE from bootstrapping was used to select hyperparameter mtry = 5).

Next, to further investigate the correlation between RNA editing and nucleotide composition and codon usages, we compared nucleotide content and specific codon usage biases between a) edited genes and transcriptome average, and b) coleoid species and nautiloids (the other cephalopod group that does not show high levels of RNA editing and is less behaviorally complex) and aplysia (a more distantly-related species). Specifically, we calculated and compared mean proportions of adenosine and GC content by gene. For codon usage, to account for the expected level of different codons, we calculated adjusted bias of a specific codon per gene as:

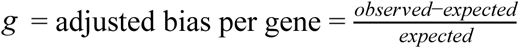

adjusted bias of a specific codon per species as:

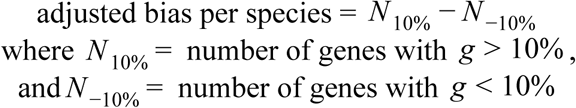

Within the four coleoid species, adjusted bias per species values were compared between edited genes (i.e. genes containing at least one editing site) and the entire transcriptome of the corresponding species as a proxy to examine the differences between edited and unedited genes. Next, adjusted bias per species values were compared between the four coleoid species and the two outgroups (nautilus and aplysia) to investigate any difference in codon usage bias that evolved during the coleoid lineage.

### VI. R/C Ratio and Amino Acid Recoding

To assess the extent of purifying selection on RNA editing in cephalopods, we calculated R/C ratios of amino acid recoding that result from editing in the four cephalopod species, as well as edits conserved in cephalopods, and edits in human, mice, and fly for comparison (using editing site data from RADAR). We classified amino acids into different groups using on 3 different metrics (from Hanada 2007 classifications A, B, and C; see Table I for an example), and counted a substitution as conservative if the change occurs within a category, and radical if the change crosses categories. For each metric, the number of radical recodings and conservative recodings are counted and their ratio is calculated. These ratios are averaged across the metrics to acquire the final R/C values for comparison.

**Table I.**
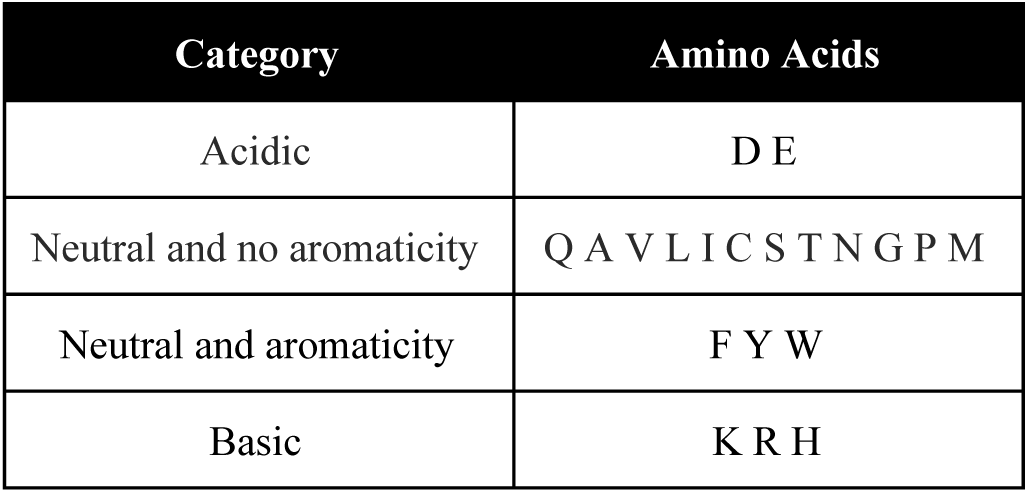
Example classification scheme of amino acids by charge and aromaticity

To gain a better understanding of the R/C results, we analyzed the distribution of specific amino acid substitutions from RNA editing. To account for inherent differences caused by the structure of the genetic code, we calculated the expected frequency of an amino acid recoding by finding the proportion of all possible amino acid changes resulting from a single A-to-I (or to G) change in a codon that cause the specific amino acid recoding. Observed frequencies of amino acid recoding by RNA editing are then compared to these expected values, and fold differences were used to compare between different amino acid substitutions and across species.

## Results

### I. Diversity Score

#### Validation of Diversity Score Metrics

To validate that the diversity score accounts for both editing level and amino acid substitutions, without one aspect being the dominating factor that determines the diversity score, we compared the diversity scores per species with (i.e. the regular diversity scores) or without (i.e. only the editing level score) the amino acid score. Since the primary function of diversity score is to identify top candidate genes, we focused the comparison on the number of overlaps in the top 100 gene list identified by scores. In general, for the 4 cephalopod species and 3 metrics (UEI, PSSM, and novelty), the proportion of overlap is approximately 50~60% (Table II), indicating that the amino acid score and the editing level each contribute to about half of the overall diversity score.

**Table II.**
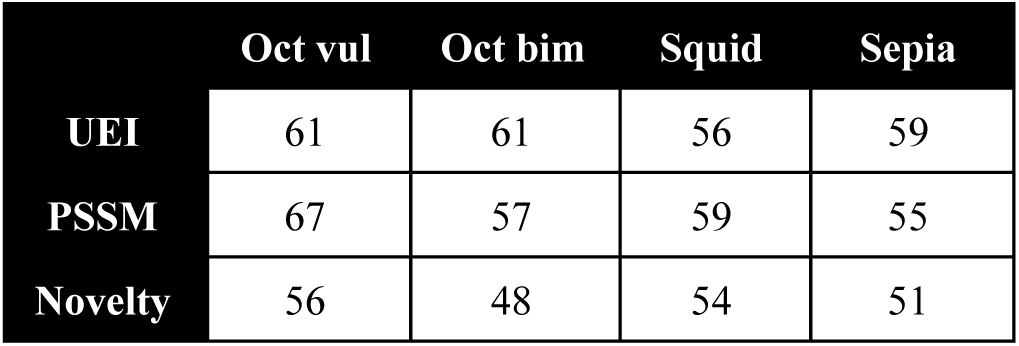
Effect of amino acid on overall diversity score metrics: each cell counts the number of overlapped genes in the top 100 gene lists identified using the specified diversity score in each species with or without the amino acid score component.

|  | Oct vul | Oct bim | Squid | Sepia |
| --- | --- | --- | --- | --- |
| UEI | 61 | 61 | 56 | 59 |
| PSSM | 67 | 57 | 59 | 55 |
| Novelty | 56 | 48 | 54 | 51 |

#### Top 100 Candidate Gene List

Table III shows the first 5 genes of the top 100 list of candidate genes as identified by the aggregate diversity score using both UEI and PSSM scores (see Supp. II for the all top 100 genes). Genes are noted for being highly connected in protein-protein interaction networks (“hub genes”), containing at least one nonsynonymous editing site in other model species (human, mice, or fruit fly), or associated with human intelligence as according to a recent GWAS study (*22*).

**Table III.** Top 5 candidate genes identified from diversity score: includes annotations from network analyses (whether if the gene is identified as a hub gene); comparison with edited genes in other species (whether if the gene contains at least one nonsynonymous mutation in human, mice, or fly); and whether the gene has been associated with human intelligence.

| Rank | Gene | Network | Conserved | Intelligence |
| --- | --- | --- | --- | --- |
| 1 | SCN9A |  |  |  |
| 2 | HUWE1 | ✓ |  |  |
| 3 | MACF1 |  |  |  |
| 4 | DOP1A |  |  |  |
| 5 | UBR4 | ✓ | Human |  |

#### Novel Editing Sites

In addition to theoretical molecular diversity that focuses on even representation of different types of transcripts, we developed another metric to identify genes with large amounts of novel editing sites. In this case, the direction of editing is important; an edit that substitutes a commonly found amino acid into a rare, novel amino acid at high editing level is taken to be highly novel, as reflected in the score.

Using this metric, the top 100 genes with the highest novelty generated were identified. Among them, 84 genes are included in the overall top 100 gene list (from combined UEI and PSSM). This overlap is expected since genes with a large number of editing sites will be ranked high in both measures.

Despite the similarities, a handful of genes were identified with high novelty scores but low overall scores. Supp. III includes top 5 genes that are ranked top 200 in the novelty list but have lower rankings in the overall list. Studying these genes at the particular editing sites with high novelty scores help understand functions of RNA editing not primarily used for creating high levels of combinatorial diversity, but focused on creating highly novel amino acid substitutions. These sites may have played a role in generating highly novel cephalopod proteins that led to significant cephalopod developments, especially in the nervous system, through evolution.

In addition to genes, the novelty score was used to identify specific editing sites that may lead to novel amino acid substitutions. Supp. IV shows the top 5 sites in each of the four cephalopod species by novelty score, and Table IV below shows the top 5 sites that are conserved across all four species. Given the high editing level, the rarity of the amino acid substitution, and the conservation across species, these editing sites may serve to generate novel, functionally significant proteins. These identified novel sites provide specific candidates for future experimental studies on the impact of RNA editing on protein functionalities.

**Table IV.** Top novel editing sites conserved across cephalopods. editing sites conserved across all 4 cephalopod species (squid, sepia, oct bim, and oct vul) in their respective proteins were ranked by the novelty score per site to identify individual novel sites. the table shows the corresponding sequence and location of the sites in oct bim.

| Rank | Gene | Oct Bim Sequence | Location in Oct Bim | Amino Acid Change |
| --- | --- | --- | --- | --- |
| 1 | CHRNA9 | comp163<br>911_c0_s<br>eq1 | 361 | Y -> C |
| 2 | VPS13A | comp166<br>111_c0_s<br>eq4 | 683 | R -> G |
| 3 | CNGA2 | comp153<br>103_c0_s<br>eq1 | 556 | R -> G |
| 4 | LRRK2 | comp166<br>025_c2_s<br>eq1 | 8914 | R -> G |
| 5 | PRICKLE2 | comp160<br>160_c0_s<br>eq1 | 591 | Y -> C |

#### Identification of Hub Genes From Protein-Protein Interaction Networks

To further investigate relationships and interactions between the candidate genes, protein-protein interaction networks of the top 100 and 200 candidate genes were produced using STRING (Fig. I). Connections inferred from experiments (protein-protein interactions), databases (KEGG pathways), co-expression, neighborhood, gene fusion, and co-occurrence were considered. Both networks have significantly more interactions than expected (PPI enrichment p-value < 1e-16), indicating that highly edited genes are involved in the same or interacting pathways.

**Figure I.**
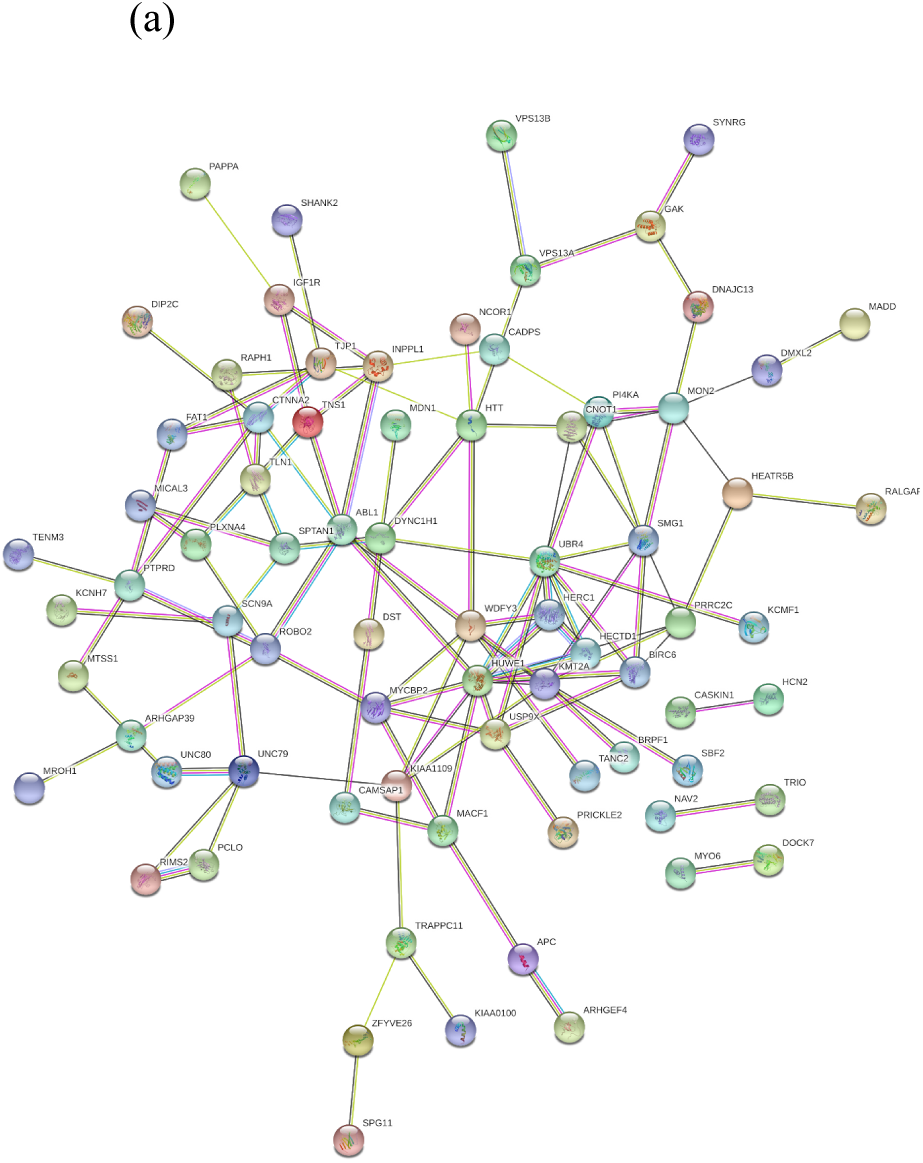
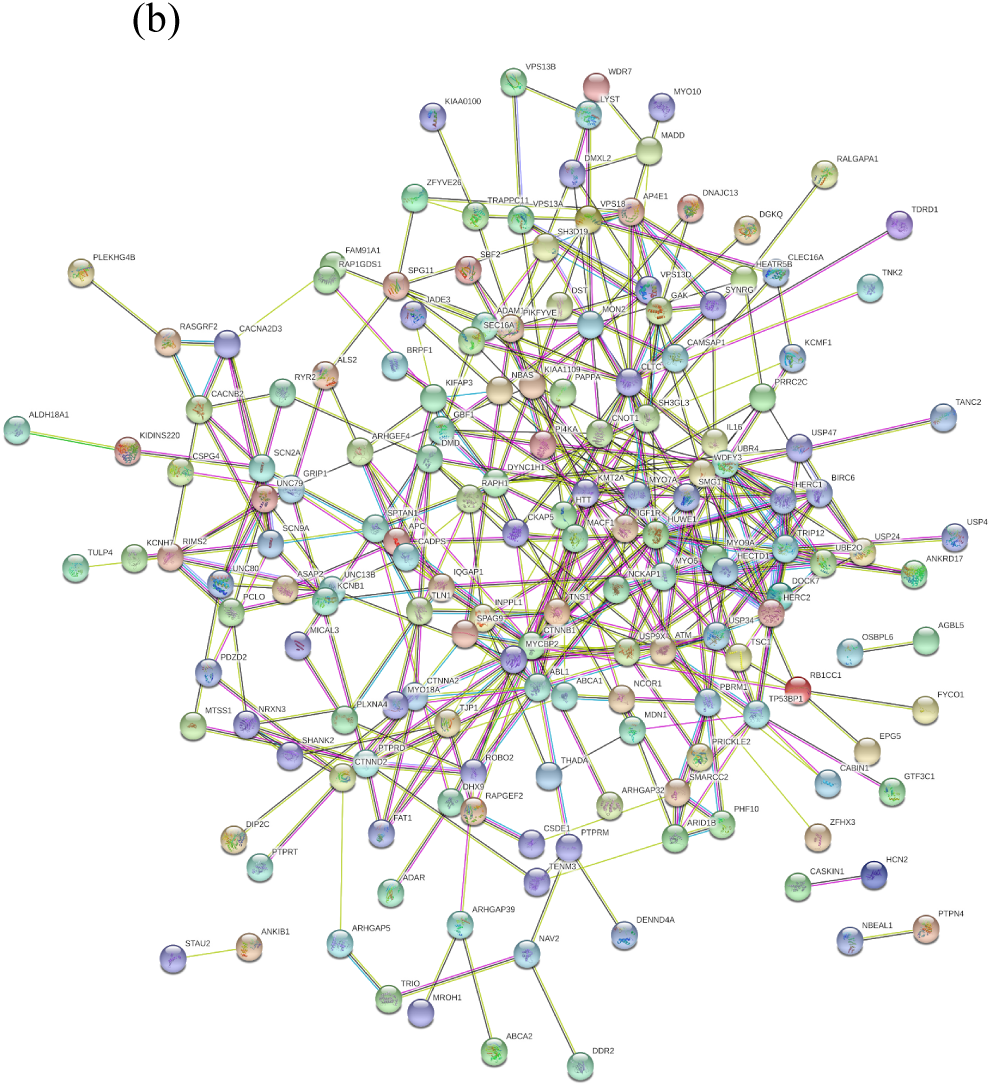
Protein-protein interaction networks of (a) top 100 and (b) top 200 candidate genes. network nodes represent proteins, edges represent interactions: known interactions from curated databases (light blue), experimentally determined (pink); predicted interactions from gene neighborhood (green), predicted gene fusions (red), gene co-occurrence (dark blue); text-mining (yellow-green), coexpression (black), protein homology (light purple). genes without any connections were removed from the graphs.

From the networks, highly connected genes we call “hub genes” were identified, which are taken to be potential key players in a diverse set of pathways and may be crucial targets of RNA editing in cephalopods. These genes are highlighted in the top 100 list (Supp. II).

#### Conservation with Model Species

Editing of top candidate genes in human, mice, and fly were investigated using data from REDIportal (*15*). In total, 11 genes were found to contain nonsynonymous editing sites in both mice and across the four cephalopod species, 22 were found for fly (*Drosophila melanogaster*) and cephalopods, and 323 were found for human and cephalopods. Comparing average diversity scores, genes also edited in mice have the highest mean score, followed by fly and human (Table V). Among the top 100 candidate gene list, 2 genes contained nonsynonymous edits in mice and fly respectively, and 20 genes contained nonsynonymous edits in human. These genes are highlighted in the overall top 100 list (Table III and Supp. II). GO analysis shows that genes edited in both cephalopods and humans are enriched in cytoskeleton-related processes, compared to genes only edited in cephalopods (Supp. V). Given the vast distance between cephalopods and these species, the conservation of editing in these genes, although not specifically by site, may indicate broad functional significance or unique structural features that facilitate their editing in a variety of species.

**Table V.** Diversity score values of various groups of genes edited in cephalopods (all edited genes, genes edited in model organisms, genes associated with human intelligence, and protocadherin-related genes)

| Cephalopod Edited Genes | Number of genes | Mean score ± SD | Mean rank ± SD |
| --- | --- | --- | --- |
| All genes | 3118 | 0.098±0.132 | 1560±900 |
| Edited in human | 323 | 0.115±0.152 | 1464±900 |
| Edited in mice | 11 | 0.263±0.237 | 641±711 |
| Edited in fly | 21 | 0.147±0.198 | 1535±1159 |
| Associated with human intelligence | 225 | 0.117±0.133 | 1380±906 |
| Clustered protocadherins | 9 | 0.065±0.028 | 1491±497 |
| Atypical cadherins | 6 | 0.419±0.285 | 199±161 |

#### Editing of Human Intelligence Genes

The candidate genes were matched to a list of genes implicated in human intelligence through GWAS and other studies (*22*). In total, 225 edited gene were identified in this intelligence set, of which 9 are in top 100 (highlighted in Table III and Supp. II). The mean score and median rank of these intelligence-implicated genes are both higher than values of the whole set (Table V), indicating a higher degree of potential diversification in these intelligence-related genes and the potential role RNA editing plays in developing complex behavior in cephalopods.

#### Editing of Protocadherin Genes

Previous studies on the octopus genome have discovered a unique expansion of protocadherins genes through gene duplication, which regulate the development of neurons and synapses (*26*). In chordates, protocadherins are also diversified using extensive alternative splicing to generate unique molecules on cell surfaces for self-recognition, a key component for the development of complex brains (*27*). In octopus protocadherins, one large exon encodes all cadherin domains, while editing is reported to be enriched in protocadherins, suggesting RNA editing, in conjunction with gene duplication, as an alternative mechanism to RNA splicing for generation of diverse protocadherin molecules in the nervous system (*27*).

To further assess the possibility that editing of protocadherins contributes to the development of neural complexity, we computed diversity scores of protocadherin genes across the four cephalopod species. As shown in Table V, edited genes in the clustered protocadherin family show a slight increase in diversity score (compared to the average diversity score across all genes), while edited genes in the atypical cadherin family do have notably higher diversity scores than expected (Table V). In fact, two atypical cadherins were ranked top 100 in the candidate list: FAT1, involved in cell migration and cell-cell adhesion, and DCHS1, which regulates neuroprogenitor cell proliferation and differentiation (*28, 29*). Interestingly, mutations in DCHS1 have been shown to cause Van Maldegem syndrome 1, an autosomal recessive disease characterized by intellectual disability and other malformations (*30*). Together, these results demonstrate the potential role of RNA editing in diversifying protocadherins and thus promoting the development of complex nervous systems.

#### Gene Ontology Analysis

Gene ontology analysis was used to determine functional enrichment of highly diversified genes. Table VI shows the 5 most significant GO terms identified from the diversity score ranked list, of which four are nervous system-related processes, indicating the diversification of RNA editing has its main effects in the nervous system. These results also reaffirm the previously reported connection between RNA editing and the nervous system in cephalopods.

**Table VI.** Top go processes of highly diversified genes and associated p-values. go enrichment and associated p-values were calculated using gorilla with a single list of genes ranked by diversity score.

| GO Term | Description | P-value |
| --- | --- | --- |
| GO:0010975 | regulation of neuron projection development | 1.88E-10 |
| GO:0042391 | regulation of membrane potential | 5.45E-10 |
| GO:0045664 | regulation of neuron differentiation | 7.92E-10 |
| GO:0051960 | regulation of nervous system development | 1.06E-09 |
| GO:0120035 | regulation of plasma membrane bounded cell projection organization | 2.01E-09 |

#### Diseases Associated with Highly Diversified Genes

Diseases enriched in highly diversified genes were investigated using DisGeNET gene-disease associations (*24*). Table VII shows top enriched diseases related to the nervous system, which include both behavioral-related disorders (e.g. autism) and molecular-based diseases (e.g. glioblastoma). These associations (with significant p-values) again highlight the connection between diversification from RNA editing and the nervous system, and also demonstrate the potential of candidate genes to aid understanding of human intelligence and related diseases.

**Table VII.** Top neural diseases enriched in highly diversified genes and associated p-values: disease enrichment was calculated using disgenet gene-disease associations dataset. p-values were calculated from the minimum hypergeometric score.

| Disease ID | Disease Name | P-value |
| --- | --- | --- |
| C0236969 | Substance-Related Disorders | 1.92E-04 |
| C0004352 | Autistic Disorder | 2.57E-04 |
| C1263846 | ADHD | 5.15E-03 |
| C0442874 | Neuropathy | 8.50E-03 |
| C0017636 | Glioblastoma | 8.80E-03 |

### II. Codon Usage Bias

#### Assessing Codon Usage Bias

Transcriptome data of four coleoid species (oct vul, oct bim, squid, and sepia) and two outgroup species (nautilus and aplysia) were used to calculate observed and expected value of all codons (except for stop codons), GC and AG content, and the codon deviation coefficient, a measure of overall level of codon usage bias, for every individual gene (*2, 12*). The observed and expected values were used to calculate adjusted bias per gene in each species, which was used to compare codon usage bias across species.

#### Predicting Editing with Nucleotide Composition and Codon Usages

To evaluate whether nucleotide composition and codon usage contains information regarding editing in general, we tested the predictive power of these information on editing in *Octopus bimaculoides*.

First, we set out to classify whether if a gene is edited using a simple logistic regression trained on genes in the transcriptome. The model achieves a decent accuracy of 81% on the test set (kappa = 0.62). Next, we attempted to predict the extent of editing (measured using cumulative editing level, normalized by length). We built a random forest model, which achieved an R-sq value of 0.23 on the test set (i.e. the model, or overall nucleotide content and codon information, can explain 23% of the variation in the extent of editing), which is not high but is expected since extent of editing depends not only on overall properties of the gene, but also specific sequence information. In fact, previous studies have been able to predict specific editing sites from SNVs with very high accuracy, but these models rely on the specific sequence surrounding the site and have only been tested on data from human and Drosophila (*31*). The predictive ability of our models, given that they only consider gene-level properties, thus indicate that indeed overall nucleotide content and codon usages are related to RNA editing.

#### Comparison of Global Nucleotide Composition

After affirming the correlation between overall nucleotide and codon information and editing, we examined the specific nucleotide composition patterns and its relationship to RNA editing in cephalopods. Given that RNA editing (specifically A-to-I editing) acts on adenosine, we expected edited genes to have higher proportions of A compared to species average, and coleoid transcriptome in general to have higher A than nautilus and aplysia. Supp. VI(a) shows the observed mean proportions of A by gene across the species: similar to our expectation, coleoids show slightly elevated proportions of A compared to nautilus and aplysia. However, edited genes do not show higher proportions of A compared to the transcriptome averages. In all cases, as expected, the range and distribution are wide. These results suggest that coleoid transcriptomes might have gone through changes that lead to elevated A, potentially facilitating RNA editing, but within each coleoid transcriptome edited genes on average maintain the same A content as other genes.

Previous studies have found regions around editing sites display higher GC content, which form stronger double-stranded regions required by ADAR enzymes for editing (*2*). Therefore, we hypothesized that edited genes would have higher GC content on average, and the widespread editing in coleoid species may have led to an increased GC content overall compared to other species. Supp. VI(b) shows the GC content per gene: contrary to our hypothesis, edited genes do not show higher GC content than all genes, and coleoid species do not have higher GC content--in fact, the median GC content is slightly elevated in nautilus, and aplysia shows significantly higher GC content than the coleoid species. Gene length does not account for this difference in GC content (R-sq < 0.01). One possible explanation is that although GC content may be higher near editing sites, other regions of the genome (e.g. crucial sites of a gene at which an edit would likely reduce functioning) may have evolved lower GC content in order to actively avoid editing.

#### Comparison of Codon Usage Bias

Given that RNA editing requires an A site and a double-stranded region near the editing site, and ADAR enzyme shows biases for nucleotides immediately upstream and downstream the A site (*2*), we hypothesized that 1) edited genes show codon usage bias that facilitate editing (e.g. for synonymous codons, the A-containing codon will be more represented to allow editing, and G-containing codon will be less represented since a codon with A can code for the same amino acid but with more flexibility), and 2) widespread RNA editing in coleoid species may have influenced codon usage in the overall genome.

To assess these potential differences, we first calculated the CDC coefficient, which measures the overall codon usage bias in a gene (*12*), and perhaps surprisingly aplysia has the largest CDC value per gene, while nautilus has about the same level of CDC as the coleoid species. Edited genes have slightly lower median CDC values, but the difference seems negligible. One possible explanation for such results is gene length: shorter genes are slightly biased towards higher CDC values (*12*), and indeed aplysia genes have shorter lengths than other species, while edited genes tend to be longer (Supp. VII), and there is a negative correlation between gene length and CDC value in the dataset (R-sq = 0.25).

Next, we analyzed biases of individual codons across the species. Comparing edited genes with the overall species as control, 23 codons are consistently more biased in edited genes for all four coleoid species, and 65% of them contain at least an A, while 48% contain G (the neutral expectation is 57%); 21 codons are consistently less biased in edited genes, of which 52% contain A, and 67% contain G. In addition, A at codon position 2 is more biased in edited genes for all four species. Together, these results supporting our hypothesis that codons with A may be biased in edited genes for allowing potential editing. Among these codons, CAG and TCT are consistently more biased by at least 10% (see Methods for details on calculations) in edited genes for all four coleoid species. As an A-containing codon, CAG codes for glutamine, which can be edited to CGG for arginine, an important type of editing that underlies a crucial RNA editing site in mice (Q/R in GRIA2, a neurotransmitter receptor that if unedited, leads to early death in mice)(*1*).

Comparing coleoid species and nautilus, the non-coleoid cephalopod with little RNA editing and less behavioral complexity, A is more biased in all four coleoid species, while G is less biased, again supporting our hypothesis that RNA editing leads to biases towards A-containing codons over G. Interestingly, C is more biased in coleoids compared to nautilus, and T is less biased, potentially to preserve similar GC content. Several A-containing codons are more biased by at least 10% in all four coleoid species compared to nautilus, including ATG, TTA, CAA, and CGA, of which two may lead to amino acid recoding through editing (ATG for M/V, CAA for Q/R). Codons AGA and AGG are less biased by at least 10%, which can be edited from AAA and AAG, a K/R amino acid change that is conservative and over-represented in cephalopods (see next section). The codon AAG and its edit to AGG also corresponds to the cephalopod ADAR preference for an upstream A and a downstream G around the editing site (*2*). Continuing these comparisons to between coleoids and aplysia, several trends still hold true: ATG, TTA, and CAA are more biased by at least 10% in all coleoids compared to aplysia as well. G at position 2 and codon AGA are more biased by at least 10% in both aplysia and nautilus compared to the coleoids.

Totaling these comparisons, codons ATG, GAA, and CAA among others are consistently more biased by 5% in 1) edited genes compared to overall species average, and 2) coleoids compared to nautilus and aplysia. A at position 2 and codon AAA are consistently more biased in the aforesaid levels, while their counterparts G at position 2 and codon AGA (corresponding to the potential edit K/R) are consistently less biased. Together, these results demonstrate the changes in codon usage bias that correspond to potential effects or causes of widespread RNA editing.

### III. Natural Selection and Amino Acid Recoding

#### Assessing Natural Selection with R/C Ratio

To assess the level of natural selection (specifically, purifying selection) on cephalopod RNA editing, R/C ratios were calculated from amino acid recoding of nonsynonymous editing sites in the cephalopod species as well as human, mice and fly for comparison. Results (Table VIII show that cephalopods have clearly lowered R/C ratios compared to human, mice, and fly, and within each group the R/C ratios are similar. This indicate that cephalopods face similar levels of purifying selection on RNA editing that are notably stronger than model species. Furthermore, edits conserved across cephalopods show even lower R/C ratio, providing evidence that conserved edits undergo stronger selection and are more likely to have some form of functional significance. Interestingly, there is no clear relationship between R/C and editing level (Supp. VIII), unlike N/S, which has been shown to increase as editing level increases, potentially because measures incorporating R/C mostly correlates with purifying selection, while N/S is also influenced by positive selection, which may be strong for edits with higher editing levels (*2, 9*).

**Table VIII.** R/C ratios of amino acid substitutions resulting from rna editing in cephalopods andmodel species

| Species | R/C |
| --- | --- |
| Cephalopod conserved | 0.52 |
| Oct bim | 0.55 |
| Oct vul | 0.57 |
| Squid | 0.58 |
| Sepia | 0.60 |
| Human | 0.79 |
| Fly | 0.79 |
| Mice | 0.83 |

Another potential explanation for the reduced level of radical amino acid substitutions from RNA editing in cephalopods is that through changes of ADAR preferences and transcriptome sequence composition, the RNA editing mechanism is biased towards potential editing sites that give rise to conservative amino acid substitutions. Since conservative substitutions are on average less likely to be deleterious, this would lower the selective cost of RNA editing and promote the evolution of widespread RNA editing (*9*).

#### Distribution of Amino Acid Recoding by RNA Editing

Given the reduced R/C ratios observed in cephalopods, we hypothesized that cephalopod RNA editing may have evolved distinct amino acid recoding preferences that favor conservative over radical substitutions, thus decreasing the potential for deleterious effects of edits. To test this hypothesis, we calculated the distribution of amino acid substitutions (measured in fold difference from expected frequencies of amino acid substitutions based on the setup of the genetic code) by RNA editing in the cephalopod species, again in comparison to human, mice, and fly. As shown in Figure IV, all four cephalopod species display similar distributions: for example, K/R and S/G substitutions are heavily over-represented in cephalopods, two conservative amino acid changes. Compared to human, fly, and mice, cephalopod amino acid recoding shows a weaker preference for radical substitutions (for example, we see reductions in K/E and E/G substitutions, both of which are radical). These results thus support our hypothesis that cephalopod RNA editing tends to result in conservative substitutions, reducing negative impacts of potential off-target edits.

**Figure IV.**
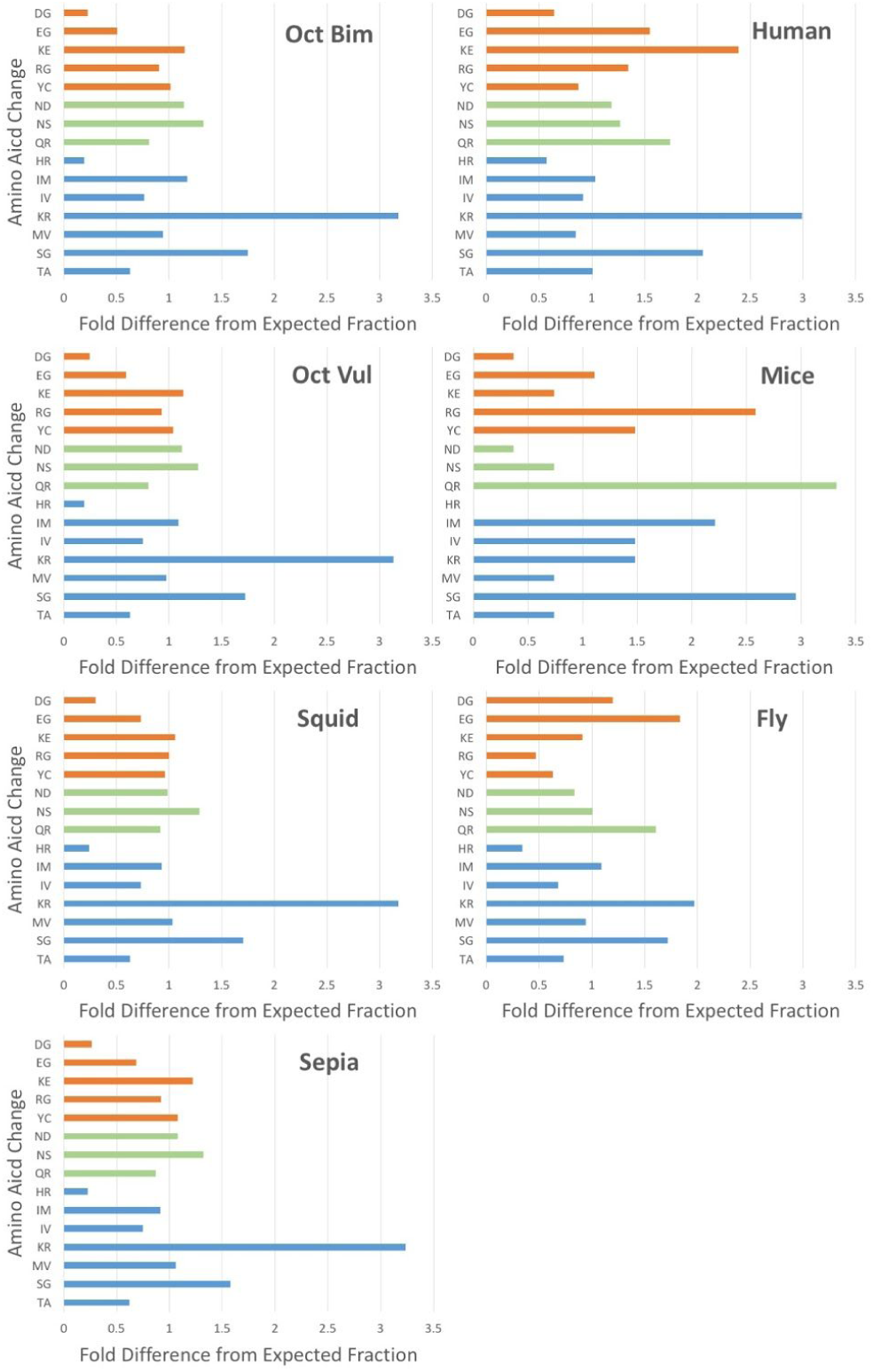
Distribution of amino acid substitutions from nonsynonymous rna editing sites in cephalopods and human, mice, and fly

## Discussion

Recently it was discovered that A-to-I editing, a rare and generally non-adaptive process in humans, leads to widespread protein recoding in soft-bodied cephalopods, a group of marine animals with large nervous systems and complex behaviors (*2*). Furthermore, neural tissues were the most highly edited and edited genes are enriched in various neural processes, suggesting a connection between RNA editing and the development of sophisticated behavior in soft-bodied cephalopods. In order to better understand the protein diversity created through RNA editing, we developed a quantitative metric that assesses the level of combinatorial diversity generated in edited proteins. This metric identifies top targets of RNA editing that may be further studied experimentally to assess the specific functional impact of the diversity brought by RNA editing. In addition, the novelty score highlights individual recoding substitutions that may be novel to cephalopods and under selective pressure, offering specific candidate sites for further investigations. As an example of experimental investigations, RNA editing on the Kv2 potassium channel orthologs in cephalopods (ranked in top 150 of the diversity score list) was recently studied by expressing the unedited and edited versions and assessing the electrical properties of the channels experimentally (*2*). Results showed that single amino acid substitutions (squid I579V, sepia I630V, and octopus I632V) lead to changes in the channels’ closing rates. Similar studies may be performed on the highlighted candidate genes and novel editing sites to help understand the molecular and cellular effects of RNA editing.

Several genes in the top 100 list may warrant further attention: ranked second in the list, the gene HUWE1 is a ubiquitin-protein ligase that mediates protein degradation (*29*). HUWE1 is also identified as a hub gene from network analyses, suggesting that it may play a central role in diverse pathways. HUWE1 is known to be involved in the regulation of neural differentiation and proliferation, and is associated to intellectual disability and severe mental retardation and developmental delays in children (Juberg-Marsidi Syndrome) (*33–35*). These connections to neural development and intelligence suggest that RNA editing of HUWE1 may play an important role in the cephalopod nervous system. Another gene of interest is PTPRD, ranked 43^rd^ in the diversity score metric, and also included in the gene set associated with human intelligence from a GWAS study (*22*). The gene codes for a protein tyrosine phosphatase that induces neuron differentiation (*29*). The gene also contains 17 editing sites conserved in all 4 cephalopod species, one of which is ranked top 5 in novel conserved edits (see Table 3). One final gene to highlight is HTT, or the Huntingtin protein, ranked 22^nd^ in the list. In humans, expansions of the polyglutamine domain in HTT lead to Huntington’s Disease, a progressive neurodegenerative disorder (*36*). HTT is also widely expressed throughout human development and is predicted to be highly pleiotropic; however, much is not known about its normal functions (*36–38*). Investigation of HTT protein diversity in cephalopods could provide a unique perspective on the function and structure of this protein.

Results of the diversity score highlight several interesting patterns and further support the connection between RNA editing and nervous system development. Disease annotations show associations of highly diversified genes with various neurological and behavioral disorders, including autism, ADHD, neuropathy, and glioblastoma. GO analyses showed highly diversified genes are implicated in neuron differentiation and neuronal projection development, a pattern noted in previous studies of RNA editing (*2*). Along with the higher levels of protein diversity in atypical cadherins and axon guidance genes (see below), these results suggest a special role for RNA editing in neuron projection development and guidance.

In addition to providing targets for future studies, these results also offer a reference dataset of protein diversity values generated by RNA editing in cephalopods. Research in cephalopod proteins may consult this data to assess the level of diversity from editing for any specific protein of interest. For example, a recent study identified the gene SLC6A4 as a receptor for drug substances that alter social behaviors in cephalopods (*39*). The level of diversity created by RNA editing on this protein can be checked using the diversity score results: in this case, SLC6A4 is edited at a negligible level, indicating that RNA editing has minimal impact on the structure and function of this protein.

Previous studies on the octopus genome have identified 15 axon guidance genes known to play an important role in the development of complex neural circuitry (*26*). Here, the diversity score data provide a perspective on the diversification of these molecules: calculations show that the mean rank of these genes in the diversity score list is significantly higher than expectation, and two genes from the set are in the top 100 list: ROBO2, a transmembrane receptor for molecular guidance cues in cell migration, and PLXNA4, a co-receptor which modulates affinity for axon growth guidance molecules (*29*). Interestingly, decreased expression of both genes were found in patients with autism (*40*). Interestingly, PLXN genes were also found to be uniquely duplicated in parrot genomes compared to all other birds, indicating PLXN gene diversity may be a more general mechanism for the evolution of advanced cognition (*41*).

### Cephalopod Intelligence: RNA editing, Autism, and Aging

Amodio et al. (2019) discuss the evolution of cephalopod intelligence and compare cephalopod biology with that of the large-brained vertebrate groups (cetaceans, corvids, primates). In contrast to the vertebrate groups, cephalopods seem to be unique in their lack of sociality. Some species aggregate into large anonymous schools, but cephalopods are generally solitary and do not engage in complex social bonds (*42*).

Similar to the asociality of cephalopods, Autism spectrum disorders in humans are characterized by dysregulation of social cognition, impaired communication, and stereotyped and repetitive behavior (*43, 44*). The high prevalence and heritability of autism spectrum disorders have led to a variety of evolutionary hypotheses on their origin and maintenance (*45*). One such hypothesis is the “solitary forager hypothesis”, which proposes that the autistic behavioral profile would have been advantageous for hunting and gathering in scarce ancestral environments (*46, 47*). Cephalopods are also characterized by a solitary lifestyle and complex foraging niche (*42*), suggesting that there may be cognitive and neural similarities between autistic humans and cephalopods. This idea is supported by the results of our disease enrichment analysis that shows autism-risk genes are highly diversified by RNA editing across four cephalopod species (see Supp. IX for further analysis). Interestingly, ADHD-related genes also show enrichment. Mental traits associated with ADHD have also been conceptualized as adaptations for the dangerous and complex foraging environments faced throughout human evolution (*46, 48, 49*).

Previous work has shown that social behavior in bees is mediated by autism-related genes (*50–52*). Taken together with our results, this indicates a common genetic toolkit that modulates sociality and foraging-related cognitive traits across both vertebrates and invertebrates. Further support for a shared genetic toolkit comes from a study showing positive selection of autism risk genes in a cave-dwelling fish that displays autistic-like traits such as asociality, hyperactivity, imbalanced attention, and repetitive behaviors (*53*).

The high degree of RNA editing in these genes in cephalopods also suggests a more specific mechanism in which a large, asocial brain can be produced. Autism-risk alleles are also known to be associated with neurogenesis and enhanced brain size, and it has been hypothesized that the widespread positive selection of these genes in humans is due to this association (*43, 54*). Positive selection on RNA editing in these genes may have played a similar role in the development of a large brain in cephalopods. In addition, evidence in humans suggests that the etiology of autism is mediated to significant degree by effects of variation in gene copy number (*55–58*). RNA editing may have played a similar role to gene duplication in creating a diversity of gene products and enhancing transcriptional complexity. Overall, this hints that modulation of a few core genetic pathways through either RNA editing or gene duplication can affect both brain size and social cognition across vertebrates and invertebrates. Support for this notion comes from recent work demonstrating dysregulation of RNA editing in autistic brains (*59*). Additionally, the autism-risk genes shown to modulate sociality in bees are enriched for RNA editing functions (*51*). Further study of the connection between RNA editing and autism in cephalopods and humans may be a promising direction for future research.

Amodio et al. (2019) also highlight the uniqueness of the cephalopod’s fast life histories (lifespans of 0.5-2 years, semelparity, absence of parental care) in contrast to the slow life histories of the large-brained vertebrate groups. They suggest that cephalopod intelligence and the fast life history evolved as a response to the loss of an external shell, which in turn led to an environment with high predation and cognitively challenging foraging niches. It is interesting to consider the connection between RNA editing, lifespan/aging, and cognition in this context. The link between RNA editing and cognition is well established on both a phylogenetic level and in specific spatiotemporal scales (*2, 60–63*). It is also apparent that RNA editing comes with a significant cost in the form of random off-target edits by the ADAR enzyme and the general potential for dysregulation. A significant amount of work has highlighted the role of RNA editing in neurodegenerative diseases and aging more generally (*59, 63–67*). We propose that this “balancing act of RNA editing” (*64*) was particularly salient during cephalopod evolution. Speaking generally, primates, corvids, and cetaceans can take their time to develop large brains and advanced cognition. Because of their long lifespans, there is also selective pressure to make a brain that is built to last. The use of RNA editing as a tool for brain development was important for these groups, but ultimately limited by the aforementioned costs. The deployment of RNA editing was largely limited to non-coding edits (*68, 69*) in order to limit the load of potentially damaging proteins with off-target recoding events. Cephalopods faced very different evolutionary pressures during their evolution. In this scenario, high predation favored the evolution of a short lifespan, but there was a concurrent pressure to rapidly develop neural and cognitive complexity. Additionally, this shift in environmental pressures caused by the loss of a shell may have happened over a relatively quick evolutionary timescale. Normal mechanisms of genome evolution (coding sequence evolution, gene duplication, expansion of non-coding regulatory DNA) may not have been able to meet this challenge of rapidly evolving a large brain that can rapidly develop over the course of a cephalopod’s short life. Widespread RNA editing enriched for protein recoding may have been the only option for developing the requisite protein diversity and transcriptional malleability (*70*). However, this came with a two-fold cost. The fact that ADAR enzymes can only edit dsRNA slowed down genome evolution and possibly constrained long-term evolutionary outcomes (*2*). The more immediate cost was the significant protein load caused by dangerous off-target protein recoding events. This protein load causes more rapid neurodegeneration and aging in cephalopods. Given the fact that there was already selective pressure to evolve a faster life history, this did not incur quite as significant of a cost and thus was evolutionary permissibly. The metaphor here would be someone who needs a house as soon as possible, but only for a short time, and therefore they build it quickly but shoddily. Research on cephalopod aging is limited, but all species studied have showed a characteristic period of senescence following sexual maturation (*71–74*). Older cephalopods show deficiencies in learning, memory, and behavior flexibility that are accompanied by neurodegeneration (*72*). Further elucidation of the neurological and molecular signatures of aging in cephalopods may yield novel insights into the evolutionary costs and benefits of widespread RNA editing and protein recoding.

### Future Directions and Conclusion

Improvement and refinement of the diversity score using additional data could yield valuable insights into the mechanisms and functions of RNA editing in cephalopod nervous systems. Generation and analysis of diversity score by specific neural tissues (the current data is pooled over multiple neural tissues) could provide insights into variations in RNA editing and protein diversity across tissues. Another direction is the acquisition of more data on RNA editing across multiple individuals for each cephalopod species. These data will help establish reliable patterns of editing and diversity and elucidate intraspecies variation in RNA editing and protein diversity. Another crucial component of the overall diversity created by RNA editing is temporal diversity, or the variation of produced proteins through time in response to varying environmental conditions or stages of development (*6*). This diversity may offer organisms greater flexibility in adapting to a changing environment but is currently not considered in the diversity score due to a lack of data on RNA editing at different time points. Acquisition of such data and the development of a temporal diversity score may help elucidate the function of RNA editing in environmental adaptation and development. In addition, the current calculations assume independence between editing sites in one RNA transcript, as the diversity score represents a theoretical maximum of the combinatorial diversity. However, a previous study has discovered substantial linkage between editing sites in *Drosophila* and cephalopods (*18*). Incorporations of these correlations between edits in one transcript may improve the diversity score to better reflect the realized protein diversity.

In summary, we develop a diversity score metric that quantitatively assesses the combinatorial protein diversity created by RNA editing in cephalopods. Results highlight connections between RNA editing and nervous system development, particularly neuron projection development and guidance, and identify highly diverse genes as candidates for future study. More generally, these results support the functional significance of RNA editing in cephalopods and suggest further research on this phenomenon could help uncover the mechanisms behind their neural and behavioral complexity.

Our results regarding nucleotide composition and codon usage biases show that they in general contain information on editing, and several specific shifts in edited genes and coleoid transcriptomes correspond to facilitating RNA editing. These changes may act together with previously shown differences in cephalopod ADAR preferences (compared to e.g. humans) for sequences near the editing site to facilitate RNA editing and also influence the resulting type of amino acid substitutions, possibly lowering the selective cost of RNA editing (*2*). Potentially, initial changes in nucleotide composition and codon usages promoted RNA editing, which in turn leads to further selective pressure for codon usage biases and changes in nucleotide composition that favor editing, creating a positive feedback loop that may have prompted the evolution of widespread RNA editing in cephalopods. The lowered R/C ratio we discovered indicates stronger purifying selection at the amino acid level, and we found distinct amino acid preferences in cephalopod RNA editing favoring conservative over radical substitutions. These changes in amino acid recoding may be a result of upstream changes in nucleotide content and codon usage biases, a connection that may warrant further inspection. Analyses of ADAR preferences and their relation to nucleotide composition, codon usage bias, and amino acid recoding in the future could test these potential scenarios and potentially help unlock the mystery of the origin of widespread, adaptive RNA editing in cephalopods.

## Supporting information

Supplemental Table 1 - Diversity Scores

## ACKNOWLEDGEMENTS

I would like to thank my mentor Mr. Erik Mohlhenrich for his guidance and encouragement throughout the project. I thank Dr. Sergey Samsonau and Mr. Joseph Li for their help on the project. I thank Dr. Joshua Rosenthal of University of Chicago and Dr. Eli Eisenberg of Tel Aviv University for their valuable advice. I thank PRISMS for providing me with the opportunity and support for research. (by M. Wang)

## Supplementary Materials

**Supp. I.**
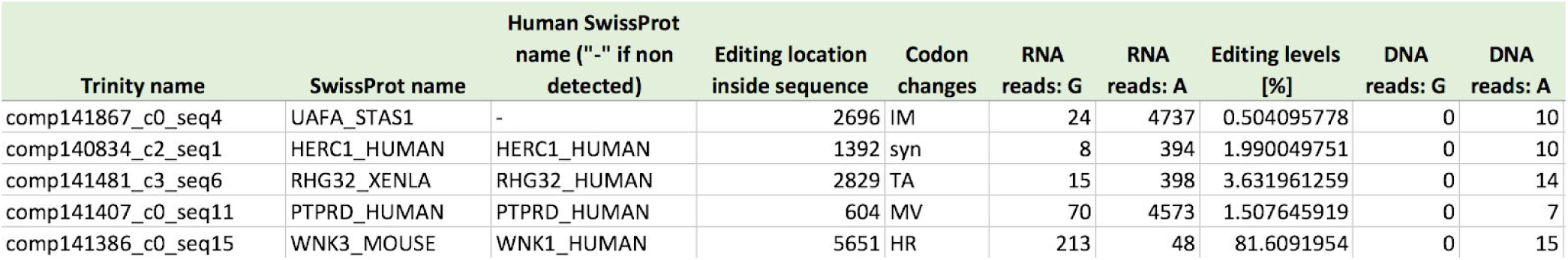
Sample rows of editing site dataset in cephalopods: trinity name, human swissprot name, editing location, codon changes, and editing levels were used in this study.

| Trinity name | SwissProt name | Human SwissProt name ("- if not detected) | Editing location inside sequence | Codon changes | RNA reads: G | RNA reads: A | Editing levels [%] | DNA reads: G | DNA reads: A |
| --- | --- | --- | --- | --- | --- | --- | --- | --- | --- |
| comp141867_c0_seq4 | UAFA_STAS1 | - | 2696 | IM | 24 | 4737 | 0.504095778 | 0 | 10 |
| comp140834_c2_seq1 | HERC1_HUMAN | HERC1_HUMAN | 1392 | syn | 8 | 394 | 1.990049751 | 0 | 10 |
| comp141481_c3_seq6 | RHG32_XENLA | RHG32_HUMAN | 2829 | TA | 15 | 398 | 3.631961259 | 0 | 14 |
| comp141407_c0_seq11 | PTPRD_HUMAN | PTPRD_HUMAN | 604 | MV | 70 | 4573 | 1.507645919 | 0 | 7 |
| comp141386_c0_seq15 | WNK3_MOUSE | WNK1_HUMAN | 5651 | HR | 213 | 48 | 81.6091954 | 0 | 15 |

**Supp. II:**
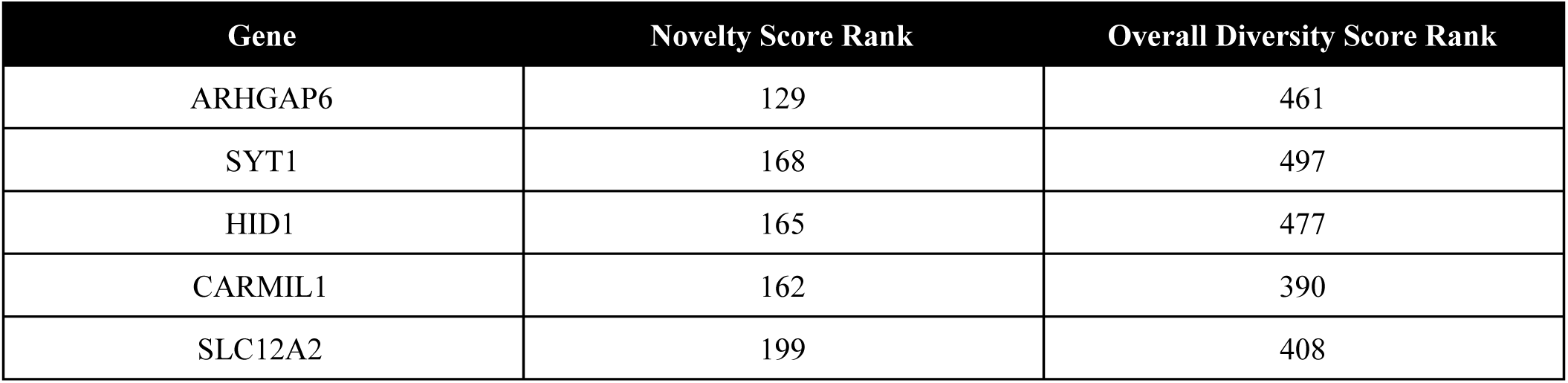
Top 100 genes by diversity score (see link)

| Gene | Novelty Score Rank | Overall Diversity Score Rank |
| --- | --- | --- |
| ARHGAP6 | 129 | 461 |
| SYT1 | 168 | 497 |
| HID1 | 165 | 477 |
| CARMIL1 | 162 | 390 |
| SLC12A2 | 199 | 408 |

**Supp. III.** Genes with highly novel editing sites not identified from overall diversity score: genes ranked top 200 in novelty scores were compared to their ranks from overall diversity score. the table displays the top 5 genes with the largest difference between the two ranks.

| Rank | Gene | Oct Bim Sequence | Location | Amino Acid |
| --- | --- | --- | --- | --- |
| 1 | ADK | comp162290_c2_seq2 | 1591 | N -> D |
| 2 | IDH3B | comp96997_c0_seq1 | 939 | N -> S |
| 3 | VPS13A | comp166353_c0_seq3 | 8736 | Y -> C |
| 4 | NPAS3 | comp164774_c0_seq1 | 1285 | Y -> C |
| 5 | MYO7A | comp164883_c0_seq19 | 5979 | S -> G |
| Rank | Gene | Oct Vul Sequence | Location | Amino Acid |
| 1 | H6PD | comp179088_c1_seq1 | 2012 | Y -> C |
| 2 | ADK | comp171017_c1_seq1 | 1061 | N -> D |
| 3 | PTPRM | comp178865_c0_seq1 | 543 | R -> G |
| 4 | IDH3B | comp178788_c0_seq1 | 1298 | N -> S |

**Supp. IV.** Top 5 novel editing sites in oct bim, oct vul, squid, and sepia.

| 5 | IGF1R | comp183541_c2_seq12 | 1153 | Y -> C |
| --- | --- | --- | --- | --- |
| Rank | Gene | Squid Sequence | Location | Amino Acid |
| 1 | ZNF570 | comp110213_c2_seq1 | 268 | H -> R |
| 2 | MYO10 | comp140327_c5_seq3 | 939 | Y -> C |
| 3 | UGGT1 | comp140978_c0_seq1 | 3070 | Y -> C |
| 4 | H6PD | comp132684_c0_seq1 | 2031 | K -> R |
| 5 | SMARCA5 | comp140138_c12_seq1 | 954 | Y -> C |
| Rank | Gene | Sepia Sequence | Location | Amino Acid |
| 1 | WNT8A | comp259845_c0_seq2 | 651 | Y -> C |
| 2 | ZNF300 | comp259368_c2_seq1 | 370 | H -> R |
| 3 | AARS | comp255397_c0_seq1 | 1063 | Y -> C |
| 4 | TOP2B | comp255128_c0_seq1 | 2720 | Y -> C |
| 5 | OTOF | comp267591_c2_seq12 | 1725 | Y -> C |

**Supp. V.** GO processes enriched in genes edited in both cephalopods and humans

SUPP. IV TOP 5 NOVEL EDITING SITES IN OCT BIM, OCT VUL, SQUID, AND SEPIA.
| GO Term | Description | P-value |
| --- | --- | --- |
| GO:0007010 | cytoskeleton organization | 1.51E-04 |
| GO:0022402 | cell cycle process | 2.90E-04 |
| GO:0031589 | cell-substrate adhesion | 5.33E-04 |
| GO:0006996 | organelle organization | 9.36E-04 |
| GO:0000910 | cytokinesis | 9.72E-04 |

**Supp. VI.**
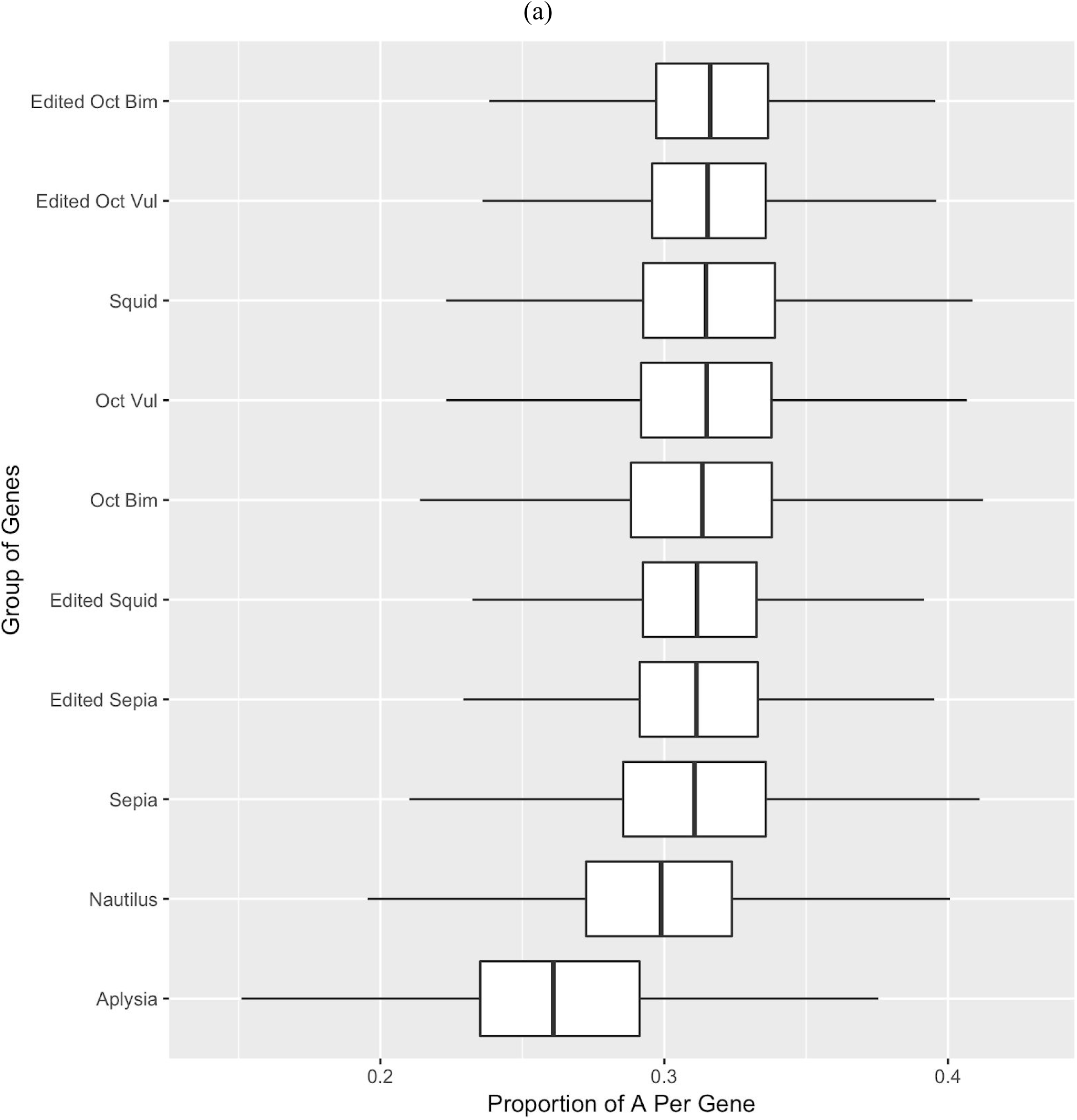
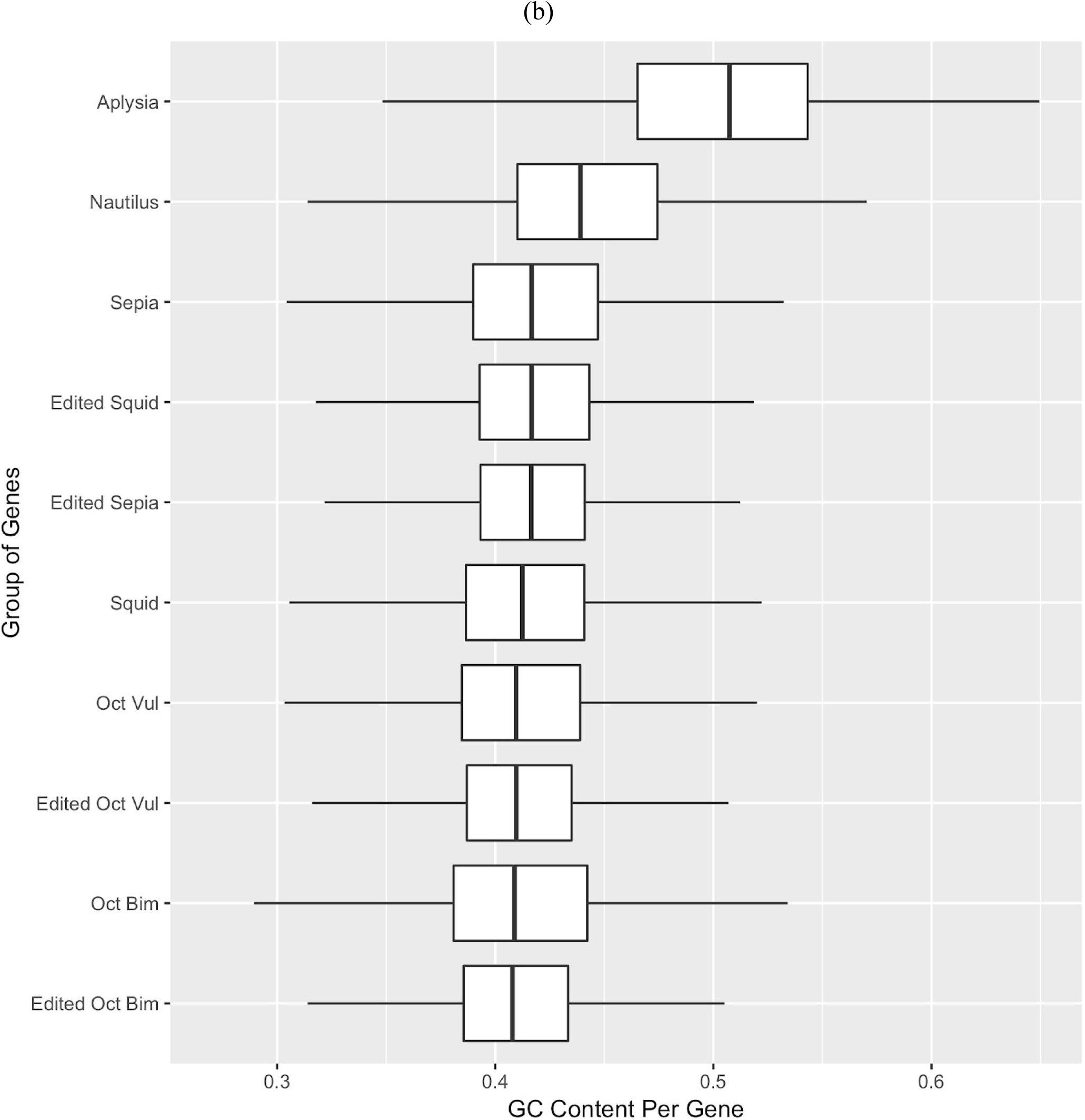
Adenosine (a) and GC content (b) per gene of edited genes, coleoid species, and outgroups. outliers are removed from the boxplot

**Supp. VII.**
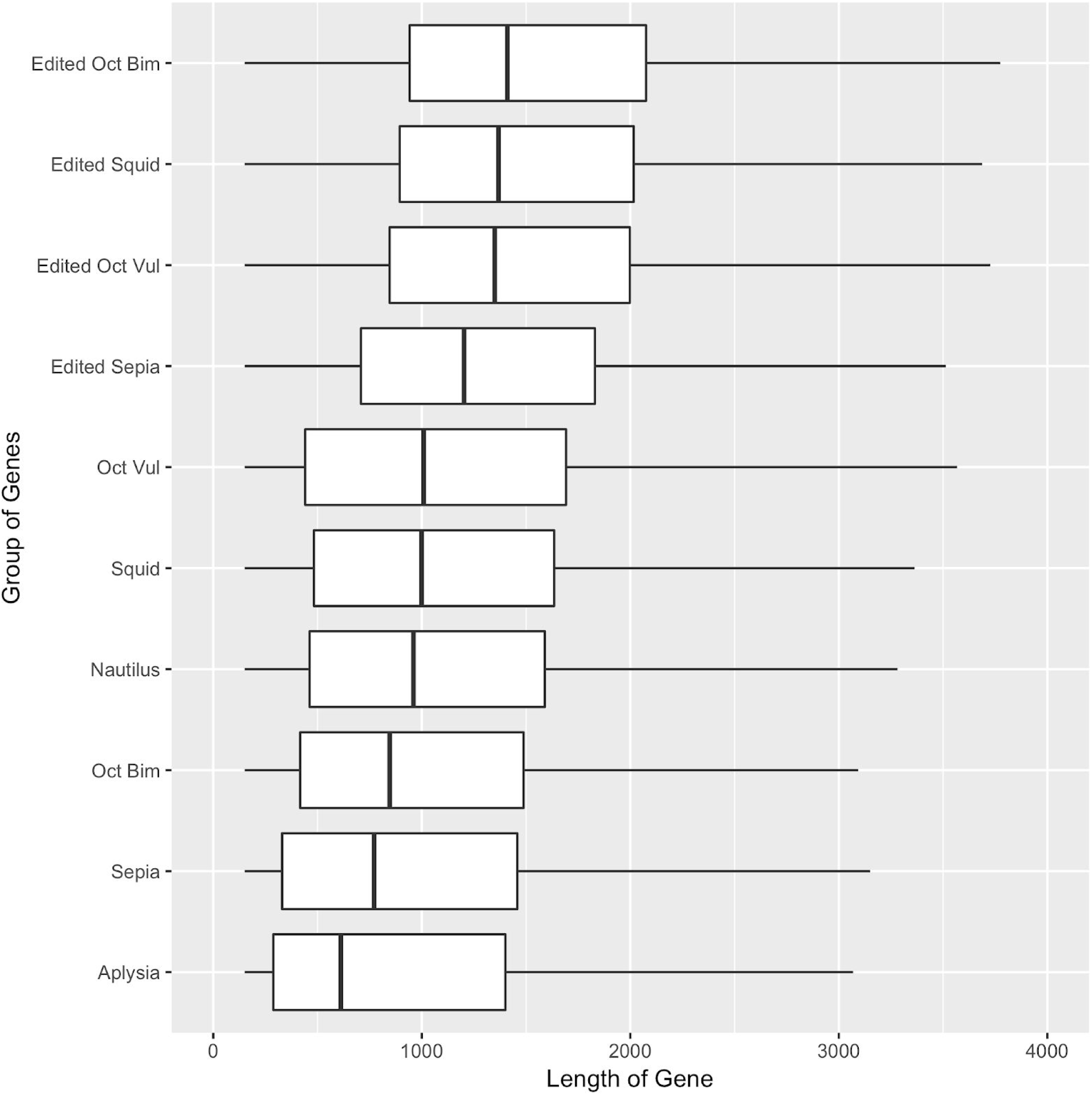
Mean gene lengths of edited genes, coleoid species, and outgroups. outliers are removed from the boxplot

**Supp. VIII.**
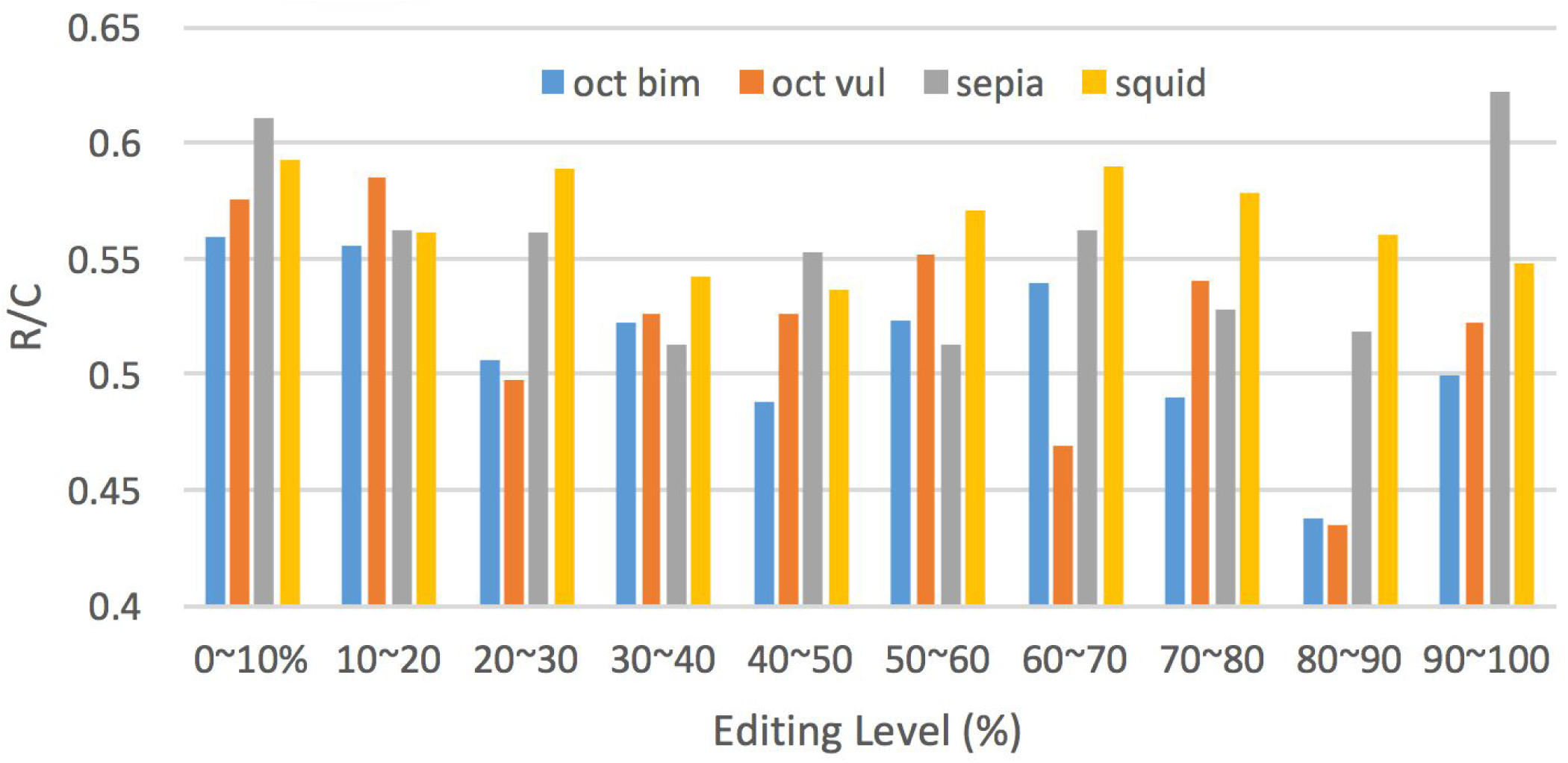
R/C ratios of editing sites by editing level in cephalopod species

**Supp. IX.** Analysis of sfari autism genes in diversity score dataset

| SFARI Autism Score | Number of genes matched in cephalopods | Average Diversity Score Rank |
| --- | --- | --- |
| 1 | 11 | 573 |
| 2 | 30 | 1015 |
| 3 | 52 | 900 |
| 4 | 106 | 1103 |
| 5 | 37 | 1470 |
| 6 | 3 | 1568 |

Further analysis of known autism genes in our diversity score dataset supports the results of the disease enrichment analysis. The SFARI database (https://gene.sfari.org/database/human-gene/) provides a list of genes connected to autism and provides a score that indicates the level of support for this connection (ranked 1-6, with 1 being the strongest evidence). The table above shows the number of genes in the SFARI database that has orthologs edited in cephalopods, broken down by the SFARI score, and their average rank by diversity score (with the neutral expectation being 1561). The elevated rank for genes with strong SFARI scores suggest that these autism-related genes are highly diversified by RNA editing in cephalopods. The top ranked gene in diversity score, SCN9A, has a score of 2 in the SFARI database. In total, 27 of the top 100 are autism-risk genes. Among them is the gene SHANK2 (SFARI score 2), ranked 82 in diversity score. The SHANK family, which organize proteins at the postsynaptic density, has been suggested as causative genes of ASD (*75*). In particular, SHANK3 is a leading autism candidate gene (score 1), with studies reporting copy number variations and points mutations in SHANK3 discovered in ASD patients (*76*). SHANK2 mutations have also been associated with ASD (*75*).

